# Aberrant type 2 dopamine receptor availability in criminal psychopathy

**DOI:** 10.1101/2023.06.21.545877

**Authors:** Lasse Lukkarinen, Jouni Tuisku, Lihua Sun, Semi Helin, Henry K. Karlsson, Niina Venetjoki, Marja Salomaa, Päivi Rautio, Jussi Hirvonen, Hannu Lauerma, Jari Tiihonen, Lauri Nummenmaa

## Abstract

**Background:** Psychopathy is characterized by antisocial behavior, poor behavioral control and lacking empathy, and structural alterations in the corresponding neural circuits. Molecular brain basis of psychopathy remains poorly characterized.

**Methods:** Here we studied type 2 dopamine receptor (D2R) and mu-opioid receptor (MOR) availability in convicted violent offenders with high psychopathic traits (n=11) and healthy matched controls (n=19) using positron emission tomography (PET). D2R were measured with radioligand [^11^C]raclopride and MORs with radioligand [^11^C]carfentanil.

**Results:** Psychopathic subjects had lowered D2R availability in caudate and putamen, and D2R striatal availability was also associated with degree of psychopathic traits in this prisoner sample. No group differences were found in MOR availability, although in the prisoner sample, psychopathic traits were negatively correlated with MOR availability amygdala and nucleus accumbens.

**Conclusions:** We conclude that D2R signaling could be the putative neuromolecular pathway for psychopathy, whereas evidence for the aberrant MOR system is more limited.

## Introduction

Psychopathy is characterized by persistent antisocial behavior, disinhibited and egotistical traits, and impaired empathy and remorse (1). It is a clinically important predictor for criminality and violence (2) and its prevalence is around 15-25% in incarcerated offenders (3). The pervasive nature of both behavioral and emotional symptoms suggest that psychopathy has organic basis, and multiple studies have found that psychopathic offenders have atrophy in frontal cortex and in limbic regions including insula and amygdala (4-8). These structural impairments are coupled with abnormal functioning of the affective division in the limbic system. Psychopathic subjects show less affect-related activity in amygdala and hippocampus, striatum and cingulate cortices while viewing emotional facial expressions, and stronger activation of frontal cortical regions (9-12). Conversely, fronto-insular functional responses are typically increased in psychopathy (12), particularly when viewing violent scenes (13). Finally, psychopaths show dampened autonomic nervous system (but typical self-evaluative) reactivity to a variety of emotional stimuli, suggesting affective disengagement (14). However, neuromolecular pathways underlying psychopathy are poorly understood.

There exist no *in vivo* imaging studies on neurotransmission in psychopathic violent offenders, yet studies on aggression and impulsivity – central aspects of the psychopathic phenotype – point towards the putative role of the endogenous dopamine (DA) and mu-opioid receptor (MOR) systems in psychopathy. DA is critical for in behavioral control, appetitive motivation, reward expectancy and prediction errors (15), and striatal dopamine system is involved in impulsive behavior in rodents (16) and humans (17). Aberrant DA functioning may predispose individuals to impulsive aggression (18), and in rats increased striatal dopamine is associated with aggressive behavior (19). However, in humans midbrain dopamine synthesis capacity is negatively associated with aggressive behavior in humans (20) and subjects with high impulsivity have lowered baseline D2R levels coupled with elevated dopamine release during reward reception (21). Finally, healthy volunteers with high psychopathic traits show amplified dopamine release during reward anticipation, indicating a link between impulsivity, psychopathy, and dopaminergic function (22).

Data from both nonhuman primates (23-25) and humans suggest that the endogenous opioid system and particularly the mu-opioid receptors (MORs) support multitude of prosocial functions including social bonding and attachment (26-29) and empathy (30, 31). Because psychopathy is consistently associated with antisocial and callous behaviour, it can be hypothesized that the psychopathic phenotype would be associated with downregulated opioidergic functioning (32, 33). However, *in vivo* imaging data on ORs in psychopathy are lacking.

### The current study

Here we tested the contribution of dopaminergic and opioidergic neuroreceptor systems to criminal psychopathy. We used *in vivo* positron emission tomography (PET) and measured availability of type 2 dopamine and mu-opioid receptors in the brains of psychopathic violent offenders and healthy control subjects. We show that psychopathy is associated with downregulated striatal D2R availability, whereas the evidence for downregulation in the MOR system is more limited.

## Methods and Materials

### Subjects

All subjects gave an informed, written consent and were compensated for their participation. The ethics board of the Hospital District of Southwest Finland approved the protocol, and the study was conducted in accordance with the Declaration of Helsinki. We studied 11 convicted violent male offenders with high psychopathic traits and 19 age and sex-matched control subjects (**Table 1)**. Offenders were inmates from the Turku Prison currently serving a sentence for murder (n=5), manslaughter (n=5), attempted manslaughter (n=3) or grievous bodily harm (n = 6). Mean recidivism rate was 2.4 times after first conviction. Prisoner subjects were screened for illicit drug use both in the screening interview and on the day of the PET scans. Details on medication and psychiatric diagnoses of the forensic subjects are presented **in Table S1**. Forensic subjects were escorted to the hospital imaging site by two prison guards who monitored the whole study protocol.

**Table 1.** Subject characteristics.

|  | **Controls**<br>Mean (Std.) | **Offenders**<br>Mean (Std.) |
| --- | --- | --- |
| **Age** | 29 (8) | 30 (6) |
| **n (males)** | 17 | 11 |
| **Education** |  |  |
| No graduation | 0 | 1 |
| Primary School | 0 | 8 |
| Second degree | 9 | 2 |
| University | 8 | 0 |
| **Psychopathy** |  |  |
| PCL-R | - | 26 (5) *n*=11 |
| LSRP primary | 22 (3) *n*=17 | 31 (4) *n*=10 |
| LSRP secondary | 13 (3) *n*=17 | 20 (5) *n*=10 |
| **Autism** |  |  |
| AQ | 11 (4) *n*=17 | 18 (7) *n*=11 |
| **Handedness** |  |  |
| Right | 14 | 9 |

Psychopathy in the convicted offenders was assessed with the Hare Psychopathy Checklist Revised (PCL-R; 3), based on semi-structured interview by an experienced forensic psychologist / psychiatrist and review of collateral information. The PCL-R measures two dimensions of psychopathic traits: primary psychopathy involving inclination to lie, lack of remorse, and callousness, and secondary psychopathy involving impulsivity, short temper and low tolerance for frustration. Psychopathic traits in the non-convicted population were measured with the Levenson Self-Report Psychopathy Scale (LSRP; 34). This self-report instrument is based on the two-factor (primary / secondary) conceptualization of the PCL. One subject had incomplete items in the secondary psychopathy scale items and was left out from the corresponding analyses.

### Data acquisition

PET imaging was carried out with GE Discovery VCT PET/CT scanner (GE Healthcare) in Turku PET center. MOR availability was measured with high-affinity agonist radioligand [^11^C]carfentanil (35) and D2R availability with high-affinity antagonist radioligand [^11^C]raclopride (36). Synthesis of [^11^C]carfentanil and [^11^C]raclopride have been described previously (37, 38). Both radioligands were administered as a rapid bolus injection, after which the radioactivity in the brain was measured for 51 minutes. Injected radioactivity doses were 251 ± 12 MBq for [^11^C]carfentanil and 259 ± 9 MBq for [^11^C]raclopride. Molar radioactivities at the time of injection and the related mass doses were 270 ± 120 MBq/nmol and 0.49 ± 0.32 µg for [^11^C]carfentanil and 290 ± 170 MBq/nmol and 0.48 ± 0.40 µg for [^11^C]raclopride. The [^11^C]carfentanil and [^11^C]raclopride PET imaging were performed on the same day > 2.5 hours apart. All PET images were reconstructed using 13 time frames (3 × 1min, 4 × 3 min, 6 × 6 min; total of 51 minutes). High-resolution anatomical T1-weighted images was obtained with 3T PET-MRI scanner (Philips Ingenuity TF PET-MR device, Philips Healthcare, Best the Netherlands) for reference and normalization purposes.

### PET preprocessing and modelling

PET images were preprocessed in MATLAB (The MathWorks, Inc., Natick, Massachusetts, United States) using Magia pipeline (39) (https://github.com/tkkarjal/magia), which utilizes SPM12 (The Wellcome Trust Centre for Neuroimaging, Institute of Neurology, University College London) in the PET data motion correction, image registration and and Freesurfer in automated ROI delineation. Six bilateral regions of interest (ROI) including amygdala, caudate, globus pallidus, nucleus accumbens, putamen, and thalamus were extracted from MRI by using FreeSurfer (http://surfer.nmr.mgh.harvard.edu). Additionally, for investigating the MOR availability, we chose six other ROIs (dorsal and rostral anterior cingulate cortices, orbitofrontal cortex, posterior cingulate cortex, insular cortex and hippocampus) that have high MOR levels. Regions for reference tissue input required in kinetic modelling (cerebellum for [^11^C]raclopride and occipital cortex for [^11^C]carfentanil) were automatically corrected for specific radioligand binding, as described previously (39).

Regional specific binding of [^11^C]raclopride and [^11^C]carfentanil was quantified as nondisplaceable binding potential (BP_ND_) using simplified reference tissue model (SRTM) (40). Parametric BP_ND_ images were also calculated for voxel level analysis with basis function implementation of SRTM (bfSRTM) with 300 basis functions. Lower and upper bounds for theta parameter were set to 0.082 1/min and 0.6 1/min for [^11^C]raclopride and 0.06 1/min and 0.6 1/min for [^11^C]carfentanil. Before the parametric image calculation, the dynamic PET images were smoothed using Gaussian kernel with 4 mm full width at half maximum to reduce the effect of noise in voxel-level bfSRTM fit. The resulting parametric images were further normalized into MNI152 space and smoothed again using Gaussian 4 mm filter.

### Statistical analysis

Univariate statistical analyses for the ROI data were performed with R software (version 3.5.1). The normality distribution of the data was evaluated with the Shapiro-Wilk test and checked visually with density and Q-Q plots. The effect of subject group on regional BP_ND_ was investigated using independent samples t test. The correlational analyses between psychopathy scores and regional radioligand availability were analyzed with Pearson’s correlation test. Full-volume analysis of the smoothed [^11^C]raclopride and [^11^C]carfentanil images were performed with SPM12 software https://www.fil.ion.ucl.ac.uk/spm/software/spm12/. The difference between prisoners and control subjects were investigated also in voxel level with Student’s t-test. Voxel-by-voxel associations between BP_ND_ and psychopathy scores (PCL and SRPS) were assessed with multiple regression in SPM12. The results were corrected for multiple comparisons by using false discovery rate (FDR) at p < 0.05.

## Results

Mean MOR and D2R availability in the groups is shown in **Figure 1**. The ROI analysis **(Figure 2** and **Table S-2)** revealed significantly lower D2R availability on the psychopaths in putamen and dorsal caudate. These effects were also confirmed in the full-volume SPM analysis (**Figure 3**). The ROI analysis (**Figure 3** and **Table S-3**) did not reveal group differences in MOR availability. The SPM analysis revealed only increased MOR availability in the prisoner group in the dorsal part of the brainstem / ventral thalamus, but this effect was not significant after correcting for multiple comparisons.

**Figure 1.**
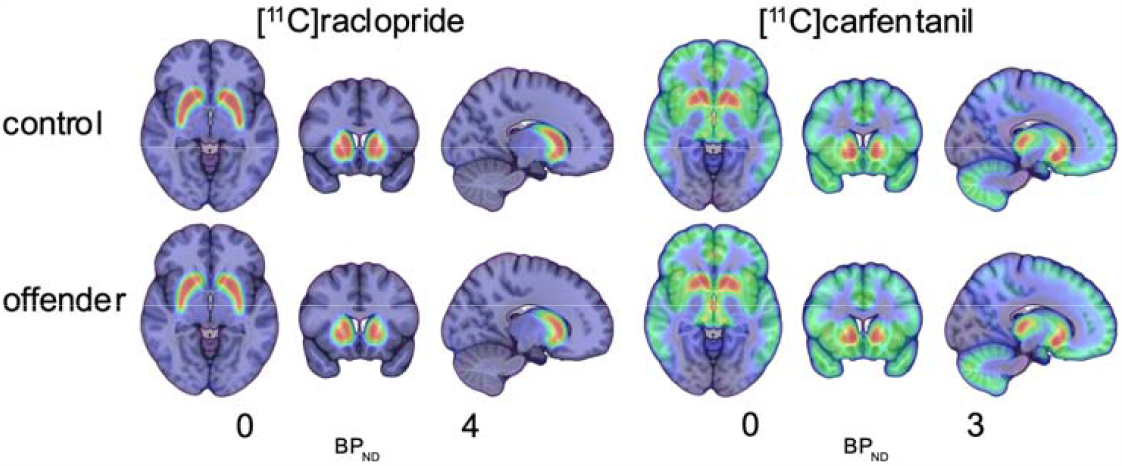
Mean D2R and MOR availability in the control and prisoner groups.

**Figure 2.**
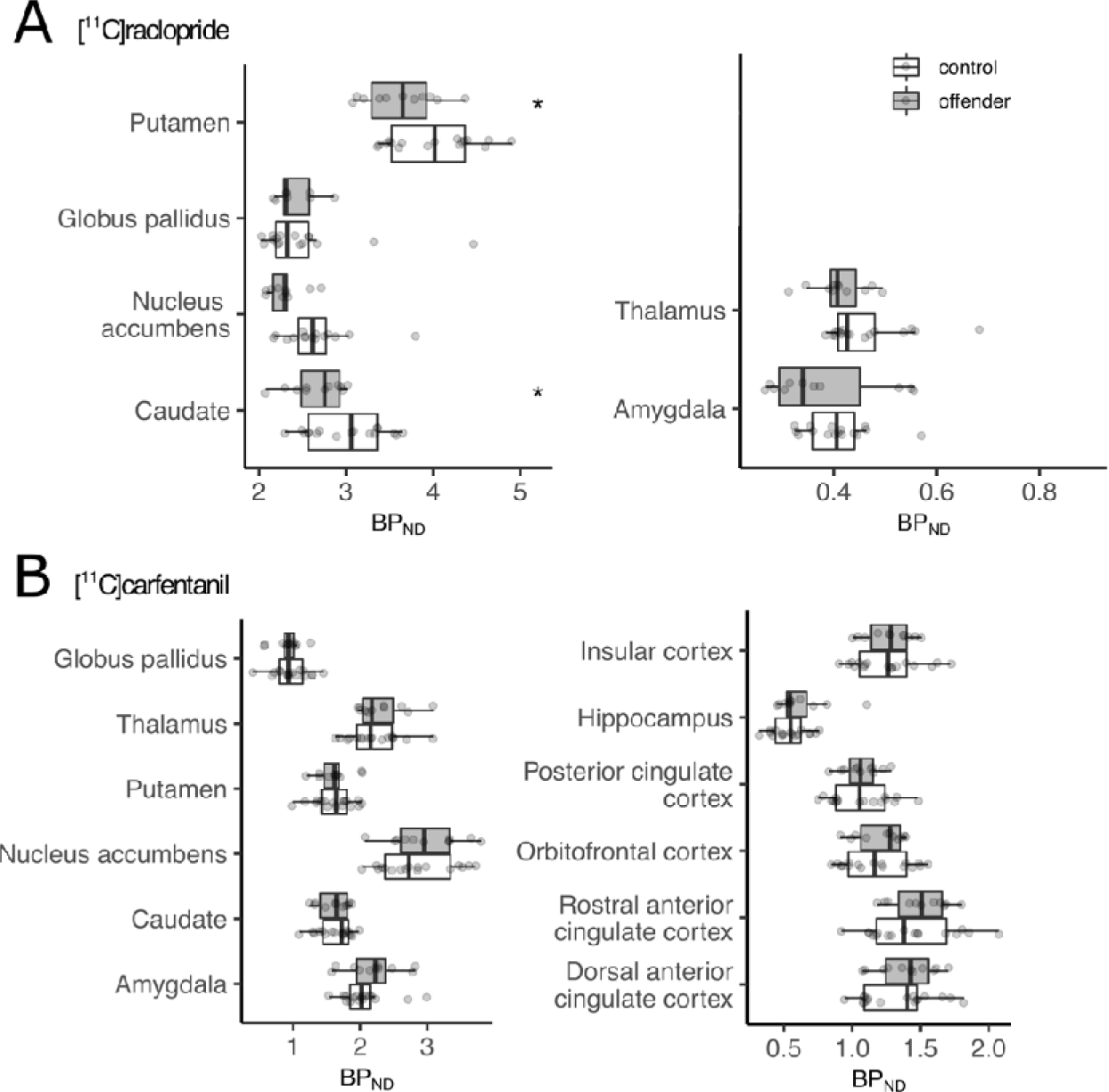
Regional group differences in (A) D2R and (B) MOR availability between control subjects and prisoners. Significant group differences are denoted with asterisk.

**Figure 3.**
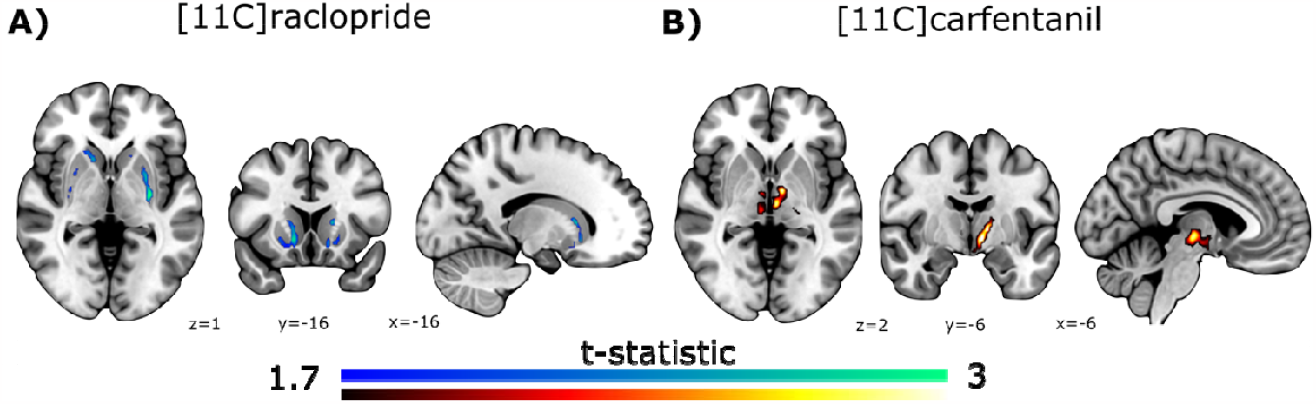
Differences between control subjects and prisoners in A) D2R and B) MOR availability. D2R results are FDR corrected at p < 0.05 (small volume correction within striatum and pallidum was utilized), whereas MOR results are shown uncorrected. Cool colours indicate regions with higher BPND in healthy controls versus prisoners, whereas hot colours indicate regions with higher BPND in prisoners versus controls.

Practically all the prisoners had history of substance abuse and there were also more smokers in the prisoners versus control group. Because both smoking and substance use may influence dopaminergic function (17, 41), the different patterns of current smoking and past drug abuse might explain the between-groups differences. To rule this possibility out, we conducted supplementary regional analysis within the offender group, where we predicted regional D2R and MOR availabilities with the psychopathy scores. This analysis (Figure 4) yielded consistently negative correlations between psychopathy and D2R availability. For the SRPS total score, the associations were significant in amygdala and nucleus accumbens; the effect for secondary psychopathy were also significant in the same areas. Similar analysis conducted with the PCL-R scores did not yield statistically significant effects.

**Figure 4.**
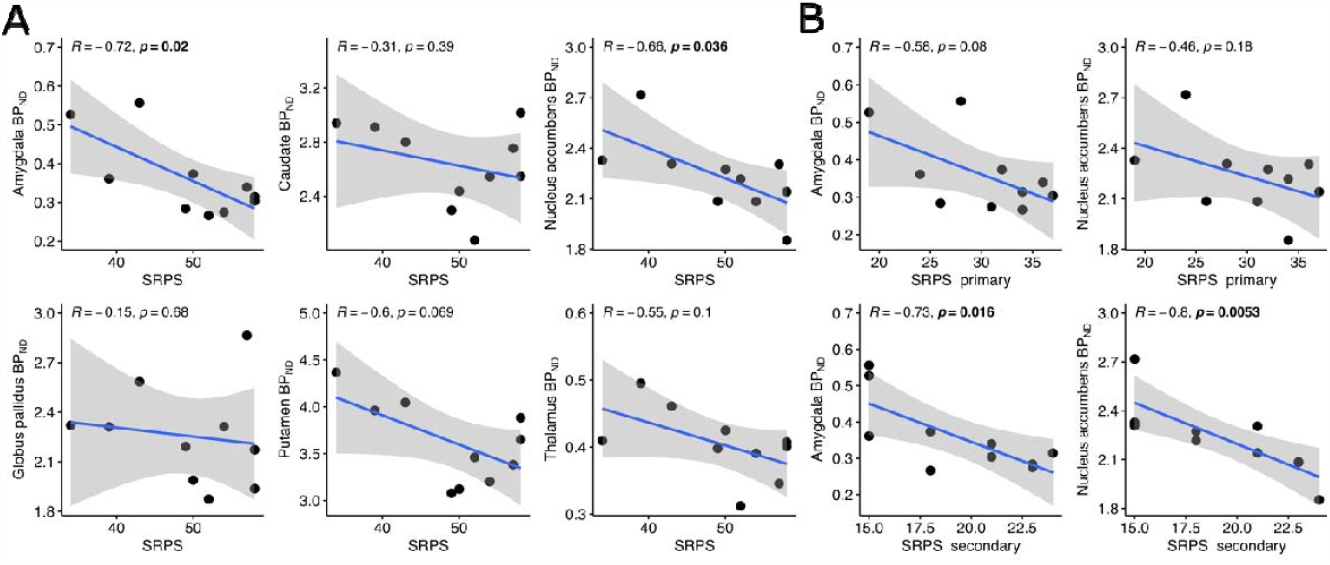
Scatterplots with LS-regression lines and 95% confidence intervals illustrating (A) the association between SRPS total scores and regional D2R availability, and (B) the association between D2R availability and primary and secondary psychopathy dimensions measured with SRPS within in the prisoner group.

Complementary voxel level correlation analysis also suggested that SRPS total scores were negatively associated with D2R availability in the striatum in the prisoner group (Figure 5A). Interestingly, in the prisoner group, PCL-R was negatively associated with MOR availability in the frontal lobe, and SRPS secondary scores were positively associated with MOR availability in thalamus (Figures 5B and 5C).

**Figure 5.**
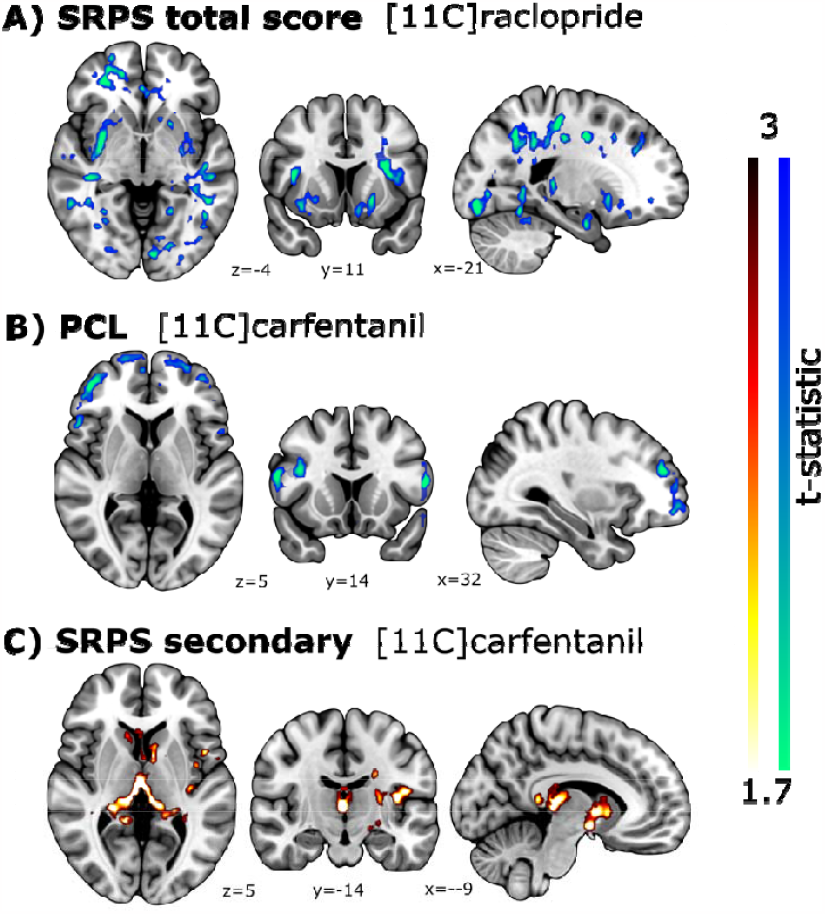
(A) Voxel level correlation between D2R availability and SRPS total scores in the prisoner group. (B) Voxel level correlation between MOR availability and PCL-R in the prisoner group. (C) Voxel level correlation between MOR availability secondary SPRS in the prisoner group. The results were corrected for multiple comparisons by using FDR at p < 0.05. Cool colors indicate a negative association and hot colors a positive association.

## Discussion

Our main finding was that psychopathy among violent offenders is associated with lowered striatal D2R availability. This effect was observed both in the direct comparison between healthy controls and subjects with psychopathy as well as correlational analysis between psychopathic traits and D2R availability in the prisoner population. There were no statistically significant differences in the MOR availability between the groups, but within-group analysis for the prisoner group revealed that MOR availability in ventral striatum and midbrain was positively correlated with psychopathic traits. These results show that aberrant D2R function is linked with criminal psychopathy, potentially explaining the aggressive and violent tendencies in this group.

### Striatal dopaminergic downregulation in criminal psychopathy

Antisocial behaviour, lack of inhibition and low empathy are all distinctive traits of psychopathy, and the first two of these traits have consistently been linked with dopaminergic neurotransmission. Dopamine system is involved in motivated behavior (42), behavioral control and particularly encoding reward prediction errors (43). Previous PET studies have shown that in noninstitutionalized subjects, low dopamine synthesis capacity as measured with FDOPA-PET is associated with impulsive/reactive aggression in laboratory tasks (44). Additionally, impulsive subjects have lowered D2R baseline availability, but exhibit increased dopamine release during reward reception (21). In line with these data, second-generation antipsychotics such as clozapine, which occupy D2R transiently are also effective in reducing aggression in addition to their antipsychotic effects (45, 46). Accordingly, the presently observed D2R dysregulation in criminal psychopathy might also reflect aggression-related traits.

Psychopaths had most consistently lowered D2R availability in striatum, a high-binding site for [^11^C]raclopride (47). Structural and functional imaging studies have however not consistently found altered structure or function of striatum in psychopathy (12, 48, 49). This likely reflects the lacking molecular specificity of these approaches, as well as the fact that the fMRI studies do not necessarily directly tap striatally mediated aspects of psychopathy. The striatal effects for lowered D2R availability however accord with the putative role of striatum in antisociality (50). In antisocial individuals, striatum may not be flexibly encoding to altered reward value and particularly lack of reward in the environment. This may explain why healthy subjects with psychopathic traits show increased dopamine release during reward (22). Accordingly, the lowered baseline D2R availability might predispose psychopaths to impulsive sensation-seeking and antisocial behavior for triggering sufficient reward signaling in the striatum.

Unlike for impulsivity and aggression, the evidence for the role of D2R in empathy and related socioemotional functions is less consistent. PET-MRI fusion imaging work has shown that specifically MOR but not D2R system is involved in empathy for pain (31). However, in monkeys, access to social contact increases D2R levels and in humans social bonding may be mediated via D2R (51, 52). Accordingly, the presently observed D2R downregulation might pertain atypical socioemotional functioning in psychopathy. In sum, the most parsimonious explaining for the present findings may be that they reflect the joint contribution of D2R to the core manifestations of psychopathy including impulsivity, aggression, and aberrant socioemotional function.

### Endogenous mu-opioid receptor system and psychopathy

Because imaging studies have consistently implicated elevated MOR tone in trait prosociality and empathy (27, 31, 53, 54) and as genetic studies also implicate the contribution of MORs in psychopathy (33) we expected that that the low trait empathy in psychopathy would link with lowered MOR availability. Against our predictions we observed no differences in MOR system tone between psychopaths and controls with the a priori statistical threshold. Weak evidence for MOR upregulation (rather than expected downregulation) in psychopathy was only found when lenient, uncorrected statistical thresholding was used. This result was however corroborated in the correlational analyses within the incarcerated sample, which confirmed that secondary psychopathic traits (pertaining to impulsivity and hostility) were positively associated with MOR availability in striatum, midbrain, and thalamus.

Because prior PET work has shown that impulsivity-related traits are associated with elevated MOR availability (55, 56), the present results may imply that MOR downregulation in psychopathy reflects psychopathy-related impulsiveness, rather than aberrant social cognition and affect. The present data thus show that associations between MOR function and extreme antisocial behaviour are nuanced. This may also reflect the complex patterns of aberrant socioemotional functioning in psychopathy. For example, psychopathic individuals show reduced frontocortical empathy responses in comparison with healthy controls (57), but this may only relate to spontaneous empathy. When deliberately asked to empathize with others, psychopaths show nearly normal responses in the brain’s social perception and empathy networks (58). In future it would thus be important to assess specifically the molecular alterations behind the different subtypes of psychopathy.

## Limitations

We did not have the information about genetic profile of the subjects and could not directly assess whether the alterations of D2R availability are genetic, or whether they pertain to lifestyle factors such as drug and alcohol abuse. We aimed at recruiting prisoner volunteers not using antipsychotics, antidepressants, or anxiolytics, yet it was impossible to recruit a fully drug-naive sample. Smoking was also more common in the prisoner sample. Although there is evidence on the acute effects of smoking on dopaminergic activity, meta-analytic data shows that the long-term effects of smoking pertain only to dopamine transporters and not D2/3R availability (41), which was measured in the present studies so this is unlikely to confound with the results. Moreover, correlational analysis within the forensic sample only also corroborated the association between psychopathy and D2R function. Finally, a single PET scan can neither establish the exact molecule-level mechanism underlying altered receptor availability cannot reveal whether the effects reflect genetic predispositions or experience-dependent plasticity in the D2R system.

## Conclusions

We conclude that criminal psychopathy is linked with downregulated D2R, whereas the contribution of MOR system is more subtle, highlighting the role of D2R in aggression and antisocial behavior in general. Together with the abnormal frontotemporal metabolism is consistently linked with violence (59-63), the lowered dopaminergic tone may contribute to impulsive, atypical sensation seeking and violence in psychopaths. Altogether these data show that dysfunctional dopamine system contributes to the psychopathic phenotype and support the idea that dopamine antagonists might be an effective treatment for violent psychopathy.

## Acknowledgements

The study was supported by the Sigreid Juselius Foundation and Academy of Finland (grants numbers 294897 and 332225, to L.N.).

