## Supplementary material for "Aberrant type 2 dopamine receptor availability in criminal psychopathy": SI file

**Table S-1.** Clinical and sociodemographic characteristics of the violent offenders.

| **Patient** | **Age** | **Diagnosis** | **Medication** | **Times of imprisonment** | **PCL-R Score** |
| --- | --- | --- | --- | --- | --- |
| 1 | 27 | SA, BP, AA, HC, | Hydroxyzine, Fluoxetine,  Quetiapine (stopped 2 days before) | 1 | 28 |
| 2 | 31 | SA, ADHD, APD, AA, HCV, APD | Levothyroxine, Amitriptyline | 1 | 29 |
| 3 | 24 | SA, PD, AA, HCV, APD | Citalopram, Buspirone, Propranolol | 1 | 36 |
| 4 | 31 | SA, ADHD, AA, APD | Hydroxyzine, Melatonin | 2 | 28 |
| 5 | 25 | SA, APD | Melatonin, Hydroxyzine,  Escitalopram*, Quetiapine*, Levomepromazine* (*stopped 3 days before) | 1 | 29 |
| 6 | 45 | None | None | 1 | 20 |
| 7 | 29 | SA, AA, APD | None | 4 | 27 |
| 8 | 39 | SA, MAD, APD | Hydroxyzine | 4 | 33 |
| 9 | 26 | SA, ADHD, HCV, APD | Hydroxyzine | 6 | 33 |
| 10 | 25 | SA, MAD, APD | Mirtazapine (stopped 1 day before) | 1 | 21 |
| 11 | 30 | SA, BP, AA, HCV, APD | Melatonin,  Buspirone*, Quetiapine* (*stopped 11 days before) | 2 | 23 |

APD=Antisocial personality disorder, ADHD=Attention-deficit hyperactivity disorder, MAD=Mood and anxiety disorder, AA=Alcohol abuse, PD=Panic disorder, SA=Substance abuse, BP=Borderline personality, HCV=Hepatitis C.

**Table S-2.** Means, standard deviations, and p-values for regional D2R availability between control subjects and prisoners in independent samples t test. Statistically significant differences are marked with an asterisk.

| **Region** | **Mean (SD) Controls** | **Mean (SD) Prisoners** | **p-value** |
| --- | --- | --- | --- |
| Amygdala | 0.40 (0.07) | 0.377 (0.11) | 0.588 |
| Caudate | 3.00 (0.44) | 2.663 (0.31) | **0.037*** |
| Globus pallidus | 2.47 (0.46) | 2.285 (0.30) | 0.357 |
| Nucleus accumbens | 2.56 (0.46) | 2.263 (0.24) | 0.056 |
| Putamen | 4.01 (0.50) | 3.63 (0.42) | **0.045*** |
| Thalamus | 0.461 (0.08) | 0.41 (0.05) | 0.078 |

**Table S-3.** Means, standard deviations, and p-values for regional MOR availability between control subjects and prisoners.

| **Region** | **Mean (SD) Controls** | **Mean (SD) Prisoners** | **p-value** |
| --- | --- | --- | --- |
| Amygdala | 2.058 (0.349) | 2.193 (0.406) | 0.38 |
| Caudate | 1.626 (0.249) | 1.611 (0.23) | 0.88 |
| Globus pallidus | 0.976 (0.259) | 0.916 (0.2) | 0.49 |
| Nucleus accumbens | 2.83 (0.535) | 2.999 (0.535) | 0.42 |
| Putamen | 1.613 (0.293) | 1.62 (0.252) | 0.94 |
| Thalamus | 2.201 (0.382) | 2.316 (0.359) | 0.43 |
| Dorsal anterior cingulate cortex | 1.335 (0.264) | 1.394 (0.211) | 0.52 |
| Rostral anterior cingulate cortex | 1.425 (0.319) | 1.499 (0.206) | 0.46 |
| Hippocampus | 0.543 (0.128) | 0.623 (0.194) | 0.24 |
| Insular cortex | 1.246 (0.247) | 1.274 (0.173) | 0.73 |
| Orbitofrontal cortex | 1.177 (0.242) | 1.213 (0.173) | 0.65 |
| Posterior cingulate cortex | 1.053 (0.222) | 1.07 (0.137) | 0.81 |
